# Xylazine does not enhance fentanyl reinforcement in rats: a behavioral economic analysis

**DOI:** 10.1101/2024.02.26.582112

**Authors:** Celsey M. St. Onge, Jeremy R. Canfield, Allison Ortiz, Jon E. Sprague, Matthew L. Banks

**Affiliations:** Department of Medicinal Chemistry, School of Pharmacy, Virginia Commonwealth University, Richmond, VA, USA 23298; Department of Pharmacology and Toxicology, School of Medicine, Virginia Commonwealth University, Richmond, VA, USA 23298; The Ohio Attorney General’s Center for the Future of Forensic Sciences, Bowling Green State University, Bowling Green, OH, USA

**Keywords:** fentanyl, xylazine, drug self-administration, behavioral economics, rat

## Abstract

The adulteration of illicit fentanyl with the alpha-2 agonist xylazine has been designated an emerging public health threat. The clinical rationale for combining fentanyl with xylazine is currently unclear, and the inability to study fentanyl/xylazine interactions in humans warrants the need for preclinical research. We studied fentanyl and xylazine pharmacodynamic and pharmacokinetic interactions in male and female rats using drug self-administration behavioral economic methods. Fentanyl, but not xylazine, functioned as a reinforcer under both fixed-ratio and progressive-ratio drug self-administration procedures. Xylazine combined with fentanyl at three fixed dose-proportion mixtures did not significantly alter fentanyl reinforcement as measured using behavioral economic analyses. Xylazine produced a proportion-dependent decrease in the behavioral economic Q_0_ endpoint compared to fentanyl alone. However, xylazine did not significantly alter fentanyl self-administration at FR1. Fentanyl and xylazine co-administration did not result in changes to pharmacokinetic endpoints. The present results demonstrate that xylazine does not enhance the addictive effects of fentanyl or alter fentanyl plasma concentrations. The premise for why illicitly manufacture fentanyl has been adulterated with xylazine remains to be determined.

## Introduction

In April 2023, the United States Office of National Drug Policy, designated illicitly manufactured fentanyl (IMF) adulterated with xylazine (“tranq”) as an emerging public threat (Gupta, 2023). From January 2019 to June 2022, the monthly percentage of IMF and xylazine-related deaths rose 276% in the United States (Kariisa et al., 2023). Xylazine is an alpha-2 agonist that is structurally similar to clonidine and routinely used as a veterinary anesthetic, but unlike clonidine, xylazine currently has no medically approved human use (Nunez et al., 2021). The combination of xylazine with IMF is colloquially referred to as “tranq dope” (Friedman et al., 2022). People who use drugs report that xylazine gives fentanyl “legs” by potentially prolonging the fentanyl “high” and decreasing injection frequency, increasing the amount of sedation, and/or delaying opioid withdrawal signs (Friedman et al., 2022; Spadaro et al., 2023).

An emerging preclinical literature suggests that xylazine combined with fentanyl enhances some fentanyl pharmacodynamic effects. For example, xylazine and other alpha-2 agonists such as clonidine and medetomidine enhance the antinociceptive potency of fentanyl in rats (Meert and De Kock, 1994). Xylazine also potentiates the hypoxic effects of fentanyl (Choi et al., 2023) and enhances fentanyl-associated lethality in rodents (Acosta-Mares et al., 2023; Smith et al., 2023). However, recent rodent studies suggest that xylazine decreases fentanyl consumption in drug self-administration procedures (Khatri et al., 2023; Sadek et al., 2024) and morphine reward in a conditioned place preference procedure (Samini et al., 2008). Whether these xylazine effects on measures of fentanyl reward and reinforcement are due to generalized sedative or cognitive impairment effects of xylazine or selective for reward/reinforcement processes are unclear. Furthermore, potential pharmacokinetic interactions that might mediate these xylazine and fentanyl pharmacodynamic interactions remain unknown. Therefore, we examined whether xylazine pharmacodynamically or pharmacokinetically enhanced fentanyl reinforcement in a rat drug self-administration model. We utilized published behavioral economic methods and analyses to rigorously determine the reinforcing effects of fentanyl alone, xylazine alone, and a range of fixed-proportion fentanyl/xylazine combinations in male and female rats (Townsend et al., 2019).

## Methods

### Subjects

A total of 23 adult Sprague-Dawley rats (11 M, 12 F) weighing 240-310 g upon arrival served as subjects (Envigo, Frederick, MD). All rats were surgically implanted with an indwelling jugular catheter and vascular access port using previously published methods (Townsend et al., 2021). Final sample sizes are reported for each experiment below and further detailed in Table S1. One female rat did not complete experiment 1 due to failed catheter patency and one female rat did not complete experiment 2 due to failed catheter patency. Animals were singly housed in a temperature and humidity-controlled vivarium and maintained on a 12-h light/dark cycle (lights off at 6:00 PM). Water and food (Teklad Rat Diet, Envigo) were provided ad-libitum in the home cage. Behavioral sessions were conducted 5–7 days per week from approximately 7 am – 11 am. Animal maintenance and research were conducted in accordance with the 2011 Guidelines of the National Institutes of Health Committee on Laboratory Animal Resources. Both enrichment and research protocols were approved by the Virginia Commonwealth University Institutional Animal Care and Use Committee.□

### Behavioral apparatus and procedures

Modular operant chambers located in sound-attenuating cubicles (Med Associates, St. Albans, VT) were equipped with retractable levers, and a set of three LED lights (red, yellow, green) mounted above each lever as previously described (Townsend et al., 2019). Intravenous (IV) drug solutions were delivered via activation of a syringe pump (PHM-100, Med Associates) located inside the sound-attenuating cubicle. After each session, intravenous catheters were flushed with 0.1 mL of a solution of cefazolin (50 mg/mL) in heparinized saline (250 U/mL). Catheter patency was verified at least every two weeks and at the conclusion of the study by IV methohexital (0.5 mg) administration and confirmation of instantaneous muscle tone loss.

### Fentanyl self-administration training

After surgical implantation of the vascular access ports, rats were trained to respond for 3.2 µg/kg/infusion IV fentanyl up to the terminal fixed-ratio (FR5) / 20-s time out schedule of reinforcement during daily 2-h sessions according to previously published methods (Townsend et al., 2019; Townsend et al., 2021).

### Experiment 1

We determined a fentanyl and xylazine dose-effect function in the drug self-administration procedure using published methods (Townsend et al., 2019). Briefly, once fentanyl self-administration training was complete in two males and seven females, saline was substituted for fentanyl on a double alternation schedule (i.e., DDSSDDSS: D, drug {3.2 µg/kg/infusion fentanyl}; S, saline). This schedule remained in effect until the number of infusions earned during the first saline session after a fentanyl session was ≤25% of the fentanyl infusions earned during that preceding fentanyl session. Upon meeting this criterion, rats began a single alternation schedule (i.e., SDSDS) until the number of saline infusions earned was ≤25% of the fentanyl infusions earned during the preceding fentanyl session for two consecutive alternations. Once these training criteria were met, we inserted test sessions into the sequence (i.e., DSTSDTDST; T, test) to determine a range of fentanyl (0.32, 1, 3.2, 10 µg/kg/infusion) or xylazine (10, 32, 100, 320 µg/kg/infusion) doses. We tested each unit fentanyl and xylazine dose once in each rat using a counterbalanced design and xylazine doses were tested up to doses that produced behavioral sedation, reduced respiratory function, or other undesirable effects. Smaller xylazine doses (0.1, 0.32, 1, 3.2 µg/kg/infusion) were first piloted in a subset of rats (for sample sizes, refer to Table S1). The xylazine dose range examined in the present study was determined based on the largest xylazine dose 320 µg/kg/infusion producing piloerection, loss of muscle tone, reduced breathing, and hypothermia at the end of the behavioral session upon visual inspection.

### Experiment 2

We determined the effects of xylazine in combination with fentanyl on drug self-administration in seven male and six female rats distinct from experiment 1 using previously published behavioral economic methods (Banks et al., 2011; Townsend et al., 2019). Briefly, we initially trained rats to self-administer 3.2 µg/kg/infusion fentanyl because this dose was the first half-log dose on the descending limb of the dose-effect function. This schedule of reinforcement was in effect until the number of infusions earned in a single session was approximately 20. Then, the FR requirement was progressively increased to FR5 and in effect until the number of earned infusions was within 20% of the running mean with no upward or downward trends for three consecutive days. The response requirement was then decreased to FR1 until the number of earned infusions reached stability criteria. The response requirement was then increased across consecutive sessions in a multi-day progressive-ratio (PR) schedule (i.e., 1, 3, 6, 10, 18, 32, 56, 100, 320, 560, 1000, 1800). This schedule remained in effect until each rat failed to complete the response requirement during a behavioral session yielding zero earned infusions. After this last session for all test drugs or drug mixtures examined, we reinstituted 3.2 μg/kg/infusion fentanyl under a FR1 schedule of reinforcement for at least two sessions and until the rat earned at least 20 infusions in a single session. 3.2 μg/kg/infusion fentanyl was tested first in all rats followed by counterbalanced determination of 10 μg/kg/infusion xylazine or saline. Three fentanyl/xylazine combinations 3.2 μg/kg/infusion fentanyl + 3.2 μg/kg/infusion xylazine, 3.2 μg /kg/infusion fentanyl + 10 μg/kg/infusion xylazine, and 3.2 μg/kg/infusion fentanyl + 32 μg/kg/infusion xylazine were also tested in the multi-day PR schedule using a counterbalanced design.

### Experiment 3

We collected blood samples from 5-6 rats (3 F, 3 M) implanted with chronic indwelling jugular venous catheters and vascular access ports as described above before drug or drug mixture administration and 15, 30, 60, 120, and 240 min following IV administration of 10 μg/kg fentanyl, 100 μg/kg xylazine, or 10 μg/kg fentanyl + 100 μg/kg xylazine. Blood was immediately transferred to K3-EDTA heparinized microcentrifuge tubes and centrifuged at 10,000 g for 15 min. The plasma supernatant was transferred to labelled storage tubes and all samples were stored at -80°C until purified and analyzed via LC-MS/MS.

### LC-MS/MS standard preparation

A stock solution was prepared by combining and then diluting fentanyl and xylazine to a concentration of 1,000 ng/mL in methanol. This solution was then further diluted to 100 ng/mL and 10 ng/mL in methanol to create stock solutions for all calibrators. The calibrators when spiked into plasma (100 μL) were at final concentrations of 100, 75, 50, 25, 10, 5, 1, and 0.5 ng/mL. The internal standard solution was prepared by diluting fentanyl-D5 and xylazine-D6 to a final concentration of 100 ng/mL in methanol. All solutions were stored in the freezer at -20°C in amber vials until use.

### LC-MS/MS plasma purification methods

Plasma was purified using protein precipitation with chilled crash solution (1% formic acid in acetonitrile). Briefly, 300 μL of crash solution, 100 μL of plasma and 30 μL of internal standard solution were aliquoted to a 1.5 mL microcentrifuge tube and placed in an Eppendorf F1.5 ThermoMixer (Eppendorf, Enfield, CT) for 10 min at a speed of 1500 rpm. The crashed plasma sample was then centrifuged for 5 min at 6,000 g. The supernatant was drawn off, transferred to HybridSPE phospholipid filters (Supelco, Bellefonte, PA) and extracted by vacuum. The remaining acetonitrile in the extractant was evaporated by a direct flow of nitrogen gas. The extractant was then reconstituted in 100 μL of water + 0.1% acetic acid and injected at an injection volume of 3 μL onto the liquid chromatography tandem mass spectrometer (LC-MS/MS).

### LC-MS/MS methods

A triple-quadrupole LCMS-8050 CL from Shimadzu U.S.A. manufacturing (Canby, OR) was utilized for sample analysis with a gradient separation method. The method utilized water + 0.1 vol% acetic acid (mobile phase A) and 100% methanol (mobile phase B) for the mobile phases and the flow rate was a constant 0.75 mL/min at 70°C over the 6.75 min gradient between 1% and 99% MeOH with a total method run time of 10 minutes. The gradient began with 1% mobile phase B until 5 minutes and then the concentration increased to 55% mobile phase B. The gradient then held until 6.76 minutes where the concentration again increased to 99% mobile phase B. This was held until 8 minutes where equilibration occurred before the next sample by decreasing mobile phase B back to 1% until the end of the 10 min run. The stationary phase consisted of a Raptor 50 mm X 2.1mm, 2.7 µm, biphenyl column (Restek, Bellefonte, PA) for the separation of the analytes and a Raptor guard column (Restek, Bellefonte, PA). Concentrations were quantified using LabSolutions Insight software (version 5.93).

### Data analysis

For experiments 1 and 2, the primary dependent measure was the number of infusions earned per 2-h session and data were plotted in GraphPad Prism (Prism 10, GraphPad, La Jolla, CA, USA). For experiment 2, data were fit using the Exponential Model of Demand (Hursh and Silberberg, 2008) via a template (available from the Institutes for Behavior Resources, http://www.ibrinc.org) then transformed to yield essential values as previously reported (Hursh and Silberberg, 2008; Townsend et al., 2019). Data for experiment 1 as well as Q_0_ and essential values determined in experiment 2 were compared using a mixed-effects analysis and the Geisser-Greenhouse correction was applied as appropriate. A significant ANOVA was followed by a Dunnet’s post hoc test as appropriate. Statistical significance was defined as p<0.05.

For experiment 3, both fentanyl and xylazine plasma concentrations versus time rectangular plots were constructed and noncompartmental analysis was conducted utilizing Phoenix WinNonlin software (Version 8.3) to estimate PK parameters. The plasma C_max_ was determined to be the highest observed plasma concentration and the corresponding timepoint was determined to be the T_max_. The terminal elimination rate constant (λz) was calculated by linear regression of the observed terminal natural log concentration versus time data and half-life (T½) was calculated as 0.693/ λz. Pharmacokinetic data are presented as the mean ± S.E.M. (except for T_max_ which is presented as the median value) for each study group. The AUC_0-_∞ represents the total area under the plasma curve from time zero to infinity and was calculated using the linear trapezoidal rule with the terminal AUC being calculated as the last measured concentration divided by λz. V_D_ is the volume of distribution of the drug and Cl_p_ is the plasma clearance of the drug and is calculated as dose/ AUC_0-_∞.

PK statistical analyses of data were performed using R (version 4.2.2). PK parameters between the fentanyl and fentanyl + xylazine group were compared using a t-test. A t-test was also used to compare PK parameters between the xylazine and xylazine + fentanyl group. Differences were considered significant at the 95% confidence level (p-value <0.05).

### Chemicals and reagents

Fentanyl HCl was supplied by the NIDA Drug Supply Program (RTI International, RTP, NC) and xylazine HCl was purchased from a commercial supplier (Aspen Veterinary Resources, Liberty, MO). Fentanyl and xylazine were dissolved in sterile saline and passed through a 0.22 µm sterile filter (Millex GV, Millipore Sigma, Burlington, MA) before IV administration. All drug doses were expressed based on the hydrochloride salt form and delivered based on individual weights as collected weekly. Cerilliant (Round Rock, TX) standard solutions of fentanyl and fentanyl-D5 were purchased from Sigma-Aldrich (St. Louis, MO). The xylazine standard solution was purchased from Cayman Chemical Company (Ann Arbor, MI). Xylazine-D6 was purchased as a powder and reconstituted at a final concentration of 100 ng/mL in methanol. Fentanyl-D5 was used as an internal standard for fentanyl and xylazine-D6 was used as an internal standard for xylazine. Lithium heparin Sprague-Dawley rat plasma that was used for the calibrators was obtained from Innovative Research (Novi, MI). Mobile phase A consisted of 0.1% acetic acid (100% LC-MS grade) purchased from Millipore Sigma (Darmstadt, Germany) and water (LC-MS grade) purchased from Fisher Chemical (Waltham, MA). Mobile phase B consisted of pure methanol (LC-MS grade) from Fisher Chemical (Waltham, MA). Formic acid (LC-MS grade) and acetonitrile (LC-MS grade) were purchased from Fisher Chemical (Waltham, MA). The compressed 5.0 ultra-high purity (UHP) grade argon gas tank used for the collision gas was purchased from Linde (Danbury, CT). The nitrogen gas tank that was used for the evaporation step was a nitrogen compressed gas tank obtained from Linde (Danbury, CT). A nitrogen generator was used to supply the heating, drying, and nebulizing gases from SouthTek (Wilmington, NC).

## Results

Figure 1A shows IV fentanyl and xylazine self-administration dose-effect functions under an FR5 schedule of reinforcement. Under these conditions, fentanyl functioned as a reinforcer at doses of 1, 3.2, and 10 μg/kg/infusion and displayed the prototypic inverted U-shaped dose-effect function (F(2.0, 26.6)=25.4, p<0.0001). In contrast, xylazine did not function as a reinforcer because no xylazine dose examined maintained behavior above saline levels up to doses that produced behavioral sedation and muscle tone loss. Based on these results, a fentanyl dose of 3.2 μg/kg/infusion and xylazine doses of 3.2, 10, and 32 μg/kg/infusion were selected for the fentanyl/xylazine combination experiments. Figure 1B shows responding for fentanyl, xylazine, saline, or a fentanyl/xylazine combination under the multi-day PR schedule. Extrapolated consumption at unconstrained price (i.e., Q_0_) was significantly lower for xylazine alone, 3.2 fentanyl / 10 xylazine, and 3.2 fentanyl / 32 xylazine compared to fentanyl alone (F(2.1,24.7)=4.8, p=0.016) (Figure 1C). However, there were no significant differences between the experimental groups when the FR value was 1. Fentanyl alone and fentanyl/xylazine mixtures, but not xylazine alone, had significantly higher essential values compared to saline (F(2.8,32.4)=15.9, p<0.0001) (Figure 1D) providing further evidence that xylazine did not function as a reinforcer in rats. There were no significant sex differences and data separated by sex are shown in Supplemental Figure 1. Figure 1E shows that xylazine (100 μg/kg IV) did not have any effect on fentanyl (10 μg/kg IV) concentrations over time. Figure 1F further displays that fentanyl (10 μg/kg IV) did not have any effect on xylazine (100 μg/kg IV) concentrations over time. The pharmacokinetic parameters assessed for either drug alone or in combination were also not significantly different (Table 1).

**Figure 1:**
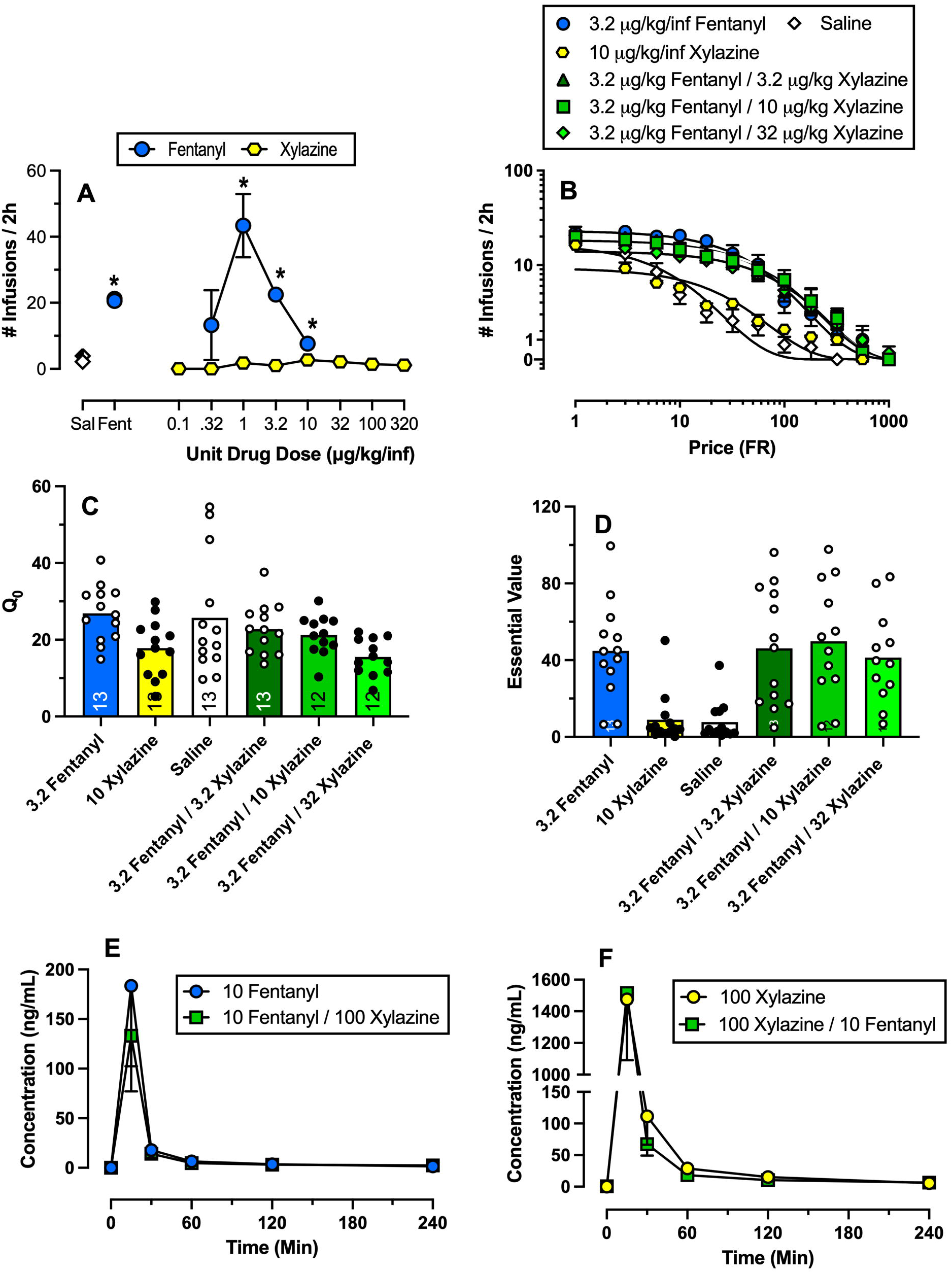
Intravenous self-administration of fentanyl, xylazine, and fixed-proportion fentanyl/xylazine combinations in male and female rats. Panel A shows fentanyl and xylazine alone self-administration as a function of unit drug dose under a fixed-ratio (FR)5 schedule of reinforcement. Points above “Sal” and “Fent” represent all training sessions before test sessions. Panel B shows fentanyl, xylazine, and saline alone or three fentanyl/xylazine fixed-proportion mixture demand curves determined in the between-session progressive-ratio drug self-administration procedure. Panels C and D are two behavioral economic metrics derived from the data in Panel B. Panels E (fentanyl) and F (xylazine) display the plasma concentration versus time values. Points in Panels A, B, E and F represent mean ± SEM whereas points in Panels C and D represent individual subjects. Asterisks in Panel A denote points that are significantly (p<0.05) different from saline. Filled points in Panels C and D denote drug or drug mixtures that are significantly different from 3.2 μg/kg/infusion fentanyl alone. For sample sizes in each experiment including distribution of females and males, refer to Table S1.

**Table 1:** Pharmacokinetic estimates for 10 μg/kg fentanyl alone, 100 μg/kg xylazine alone, and a 10 μg/kg/ 100 μg/kg fentanyl/xylazine combination. Data are mean ± SEM or median of 3-6 rats per group.

| | Fentanyl | Fentanyl<sup>\$</sup> +<br>Xylazine | Xylazine | Xylazine<sup>@@</sup> +<br>Fentanyl |
| --- | --- | --- | --- | --- |
| T<sub>1/2</sub> elimination (h) | 1.99 ± 0.38 | 2.55 ± 0.49 | 1.25 ± 0.15 | 1.61 ± 0.31 |
| AUC<sub>0-∞</sub> (ng/mL × h) | 74.11 ± 32.13 | 61.17 ± 23.35 | 472.31 ± 206.00 | 464.25 ± 181.45 |
| C<sub>max</sub> (ng/mL) | 183.45 ± 81.11 | 133.15 ± 56.01 | 1476.03 ± 729.77 | 1515.15 ± 652.60 |
| T<sub>max</sub> (h) (median) | 0.25 | 0.25 | 0.25 | 0.25 |
| Cl<sub>p</sub> (mL/h) | 129.71 ± 98.53 | 157.00 ± 81.77 | 225.82 ± 93.39 | 286.10 ± 185.67 |
| V<sub>D</sub> (mL) | 390.48 ± 302.87 | 503.85 ± 245.86 | 333.77 ± 109.51 | 534.16 ± 284.16 |

<sup>\$</sup>-Indicates Fentanyl pharmacokinetic parameters for fentanyl/xylazine mixture.
<sup>@@</sup>-Indicates Xylazine pharmacokinetic parameters for fentanyl/xylazine mixture.

## Discussion

The present study determined pharmacodynamic and pharmacokinetic effects of fentanyl and xylazine alone and in fixed-proportion combinations in female and male rats. There were two main findings from this study. First, xylazine fixed-proportion combinations with fentanyl failed to significantly alter fentanyl reinforcement using behavioral economic metrics. These results suggest that xylazine did not enhance the addictive effects of fentanyl. Second, fentanyl and xylazine pharmacokinetic parameters were similar when determined alone and in combination. These results suggest that fentanyl and xylazine pharmacodynamic interactions are not due to pharmacokinetic factors such as increased plasma drug levels. Although the precise reasons for clinical adulteration of fentanyl with xylazine remain to be empirically elucidated, these results do not support xylazine-induced enhancement of fentanyl reinforcement as a potential mechanism.

Fentanyl functioned as a reinforcer in male and female rats. The present fentanyl results are consistent with the extant intravenous drug self-administration literature in rats (Seaman and Collins, 2021; Townsend et al., 2019) and nonhuman primates (Berro et al., 2022; Maguire and France, 2022). In contrast to fentanyl, xylazine did not function as a reinforcer under an FR5 schedule of reinforcement up to doses that produced sedation, loss of muscle tone, and hypothermia (experiment 1). In addition, xylazine essential value was not significantly different from saline under the between-session PR procedure (experiment 2) providing additional evidence that xylazine did not function as a reinforcer. Consistent with the present results, xylazine failed to produce a rewarding effect using conditioned place preference in male mice (Acosta-Mares et al., 2023). However, another alpha-2 agonist, clonidine, was self-administered above vehicle levels in baboons (Weerts and Griffiths, 1999) and rats (Shearman et al., 1981), but was not self-administered more than placebo in opioid-dependent humans (Preston et al., 1985). Interactions between fentanyl and different alpha-2 agonists such as clonidine or dexmedetomidine on reinforcement endpoints remain to be fully elucidated. Overall, the fentanyl and xylazine alone results in these two drug self-administration procedures provide the empirical foundation to determine subsequent fentanyl and xylazine pharmacodynamic interactions.

Using a behavioral economic framework, increasing fixed-proportions of xylazine combined with fentanyl resulted in a selective xylazine proportion-dependent *decrease* in extrapolated fentanyl/xylazine consumption at unconstrained price without significantly altering reinforcement measures such as essential value. However, increasing fixed proportions of xylazine combined with fentanyl did not significantly attenuate responding at FR1 suggesting that fentanyl consumption was unaffected by xylazine up to the largest doses tested. These findings are generally consistent with and expand upon previous rat drug self-administration studies suggesting that xylazine decreases fentanyl consumption when combined or when xylazine was administered before a fentanyl self-administration session (Khatri et al., 2023; Sadek et al., 2024). Other studies have shown that xylazine decreased morphine reward expression in conditioned place preference but had no effect on fentanyl conditioned place preference (Acosta-Mares et al., 2023; Samini et al., 2008). In summary, the accumulating preclinical evidence suggests xylazine does not enhances the addictive effects of fentanyl.

In contrast to reinforcing effects, an emerging literature suggests that fentanyl and xylazine combinations might exacerbate undesirable effects. For example, preclinical studies suggest that xylazine enhances brain hypoxic and lethal effects of fentanyl across a broad range of experimental endpoints (Acosta-Mares et al., 2023; Choi et al., 2023; Smith et al., 2023). Thus, if xylazine does not enhance the addictive effects of fentanyl and people who use drugs describe xylazine as an undesirable adulterant (Friedman et al., 2022; Spadaro et al., 2023), why would clandestine chemists or dealers adulterate IMF with xylazine? Anecdotal reports suggest that xylazine might reduce the frequency of IMF use (Friedman et al., 2022; Rubin, 2023). Visual observations of rats self-administering a fentanyl/xylazine mixture at FR1 included muscle tone loss at the end of the first couple of behavioral sessions; however, tolerance developed quickly to these sedative effects as drug self-administration sessions continued. These observations provide some evidence to support these anecdotal reports that xylazine might reduce IMF use frequency, at least initially, due to behavioral sedative effects.

In conclusion, we offer a speculative hypothesis regarding the IMF adulteration with xylazine. One insidious hypothesis is that xylazine is being added to IMF not to enhance the addictive effects of fentanyl, but as a vector for use as a chemical weapon by hostile groups that warrants further research and investigation (Pitschmann and Hon, 2023). Support for this speculative hypothesis comes from the United States Prohibition Era where ethanol was intentionally adulterated and has parallels to the current opioid crisis where increased potency and product contamination exacerbated mortality rates (Greenwood et al., 2022).

## Notes

### Competing Interest Statement

The authors have declared no competing interest.

### Summary of Updates

Revised to update clarification in methodology and interpretation of results.

