## Supplemental Table and Figure 1 for "Xylazine does not enhance fentanyl reinforcement in rats: a behavioral economic analysis"

**Running title:** Fentanyl-Xylazine interactions in rats

^*^ Both authors contributed equally

^#^ Corresponding authors

**Table S1.** Sample sizes given by experiment and treatment.

| **Experiment** | **Treatment (µg/kg/inf)** | **n** | |
| --- | --- | --- | --- |
|  |  | **Male** | **Female** |
| **1** | 0.32 Fentanyl | 2 | 6 |
|  | 1 Fentanyl | 2 | 6 |
|  | 3.2 Fentanyl | 2 | 6 |
|  | 10 Fentanyl | 2 | 6 |
|  | 0.1 Xylazine | 0 | 1 |
|  | 0.32 Xylazine | 0 | 1 |
|  | 1 Xylazine | 1 | 3 |
|  | 3.2 Xylazine | 1 | 1 |
|  | 10 Xylazine | 2 | 5 |
|  | 32 Xylazine | 2 | 6 |
|  | 100 Xylazine | 2 | 5 |
|  | 320 Xylazine | 2 | 6 |
| **2** | 3.2 Fentanyl | 7 | 6 |
|  | 10 Xylazine | 7 | 6 |
|  | Saline | 7 | 6 |
|  | 3.2 Fentanyl/ 3.2 Xylazine | 7 | 6 |
|  | 3.2 Fentanyl/ 10 Xylazine | 7 | 5 |
|  | 3.2 Fentanyl/ 32 Xylazine | 7 | 5 |
| **3** | 10 Fentanyl | 3 | 3 |
|  | 100 Xylaxine | 3 | 3 |
|  | 10 Fentanyl/ 100 Xylazine | 4 | 1 |

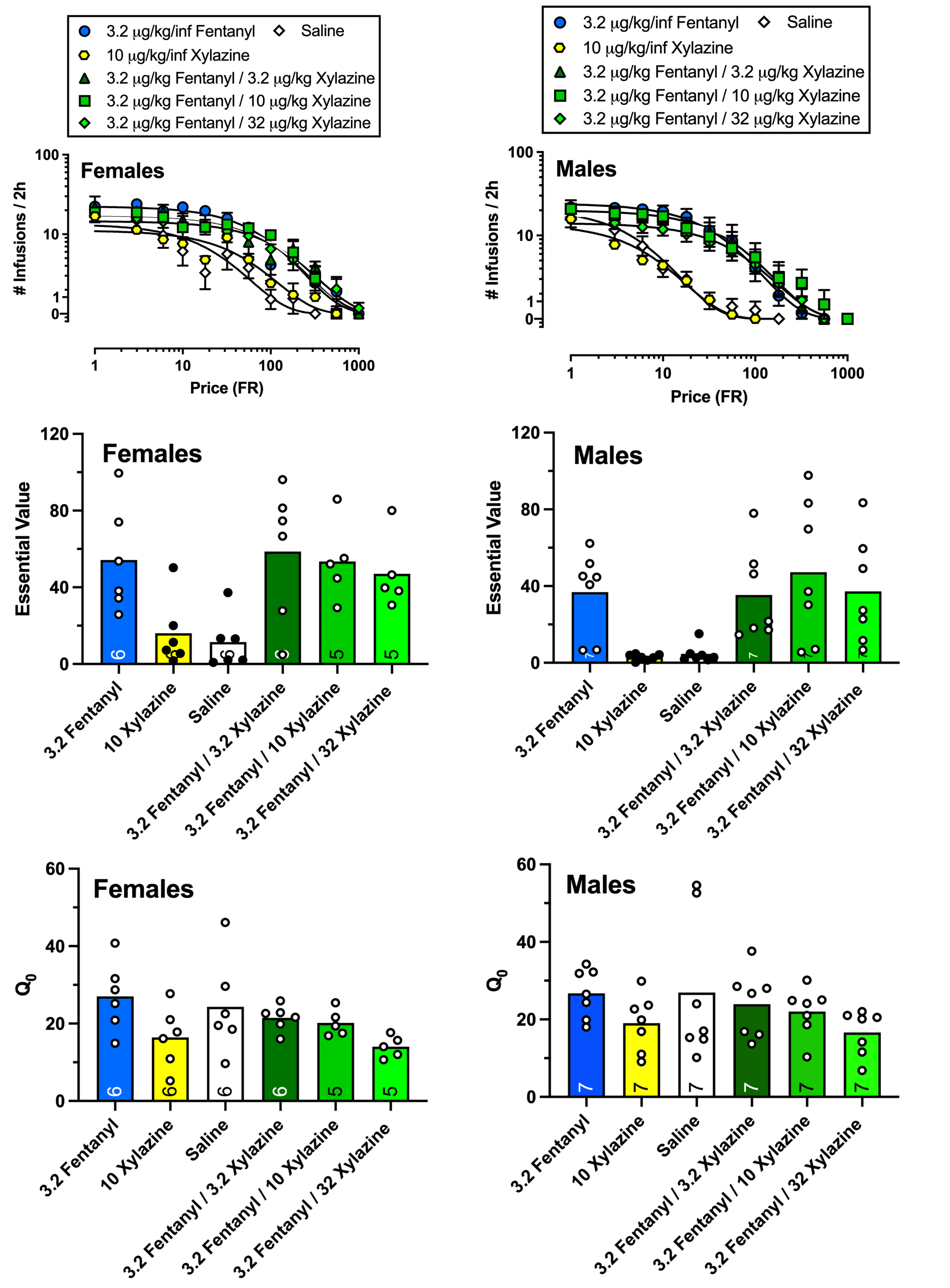

**Figure S1:** Intravenous self-administration of fentanyl, xylazine, and fixed-proportion fentanyl/xylazine combinations in female (left column) and male (right column) rats. Top Panels show fentanyl, xylazine, and saline alone or three fentanyl/xylazine fixed-proportion mixture demand curves determined in the between-session progressive-ratio drug self-administration procedure. Middle and bottom panels are two behavioral economic metrics derived from the data in top panels. Panels E (fentanyl) and F (xylazine) display the plasma concentration versus time values. Points in top panels represent mean ± SEM whereas points in middle and bottom panels represent individual subjects. Filled points in middle panels denote drug or drug mixtures that are significantly different from 3.2 μg/kg/infusion fentanyl alone (male EV: F(2.59,15.56)=7.53, p=0.0032; female EV: F(1.17,5.39)=8.4, p=0.0288).
